# ComBat-met: Adjusting Batch Effects in DNA Methylation Data

**DOI:** 10.1101/2024.08.13.607838

**Authors:** Junmin Wang

## Abstract

Integration of genomics data is routinely hindered by unwanted technical variations known as batch effects. Despite wide availability, existing batch correction methods often fall short in capturing the unique characteristics of DNA methylation data. We present ComBat-met, a beta regression framework to adjust batch effects in DNA methylation studies. Our method fits beta regression models to the data, calculates batch-free distributions, and maps the quantiles of the estimated distributions to their batch-free counterparts. Compared to traditional methods, ComBat-met followed by differential methylation analysis shows improved statistical power without compromising false positive rates based on simulated data. Additionally, we demonstrate the ability of ComBat-met to remove cross-batch variations and recover biological signals using data from The Cancer Genome Atlas.

## 1 Introduction

High-throughput genomics technologies have revolutionized our ability to investigate complex biological systems. Among these, DNA methylation profiling has emerged as a powerful tool for studying epigenetic regulation, providing insights into gene expression, organoid development, disease mechanisms, and environmental influences.

A key step in DNA methylation profiling, whether through bisulfite sequencing or methylation microarrays, is the conversion of unmethylated cytosines to thymines via bisulfite treatment (Frommer et al. 1992). This process is crucial for distinguishing between methylated and unmethylated cytosines, as only unmethylated cytosines are converted, leaving methylated cytosines unchanged. However, variations in the efficiency of this cytosine to thymine conversion across experimental batches can introduce systematic biases in methylation data (Ross et al. 2022; Price and Robinson 2018). Factors such as differences in bisulfite treatment conditions, variations in DNA quality or quantity, and technical inconsistencies in the bisulfite conversion reaction itself can all lead to batch effects (Olova et al. 2018). Failure to address batch effects can obscure true biological signals, impeding the accuracy and reproducibility of downstream analyses (Wang and Novick 2024; Akulenko, Merl, and Helms 2016).

Various methods have been developed to correct for batch effects in genomics data. Among them, ComBat, which uses an empirical Bayes framework by borrowing information across genes, has gained widespread adoption in the field of microarray data analysis (Johnson, Li, and Rabinovic 2007). ComBat-seq, which employs a negative binomial regression model to retain the integer nature of sequence count data, is widely used in RNA-seq studies (Zhang, Parmigiani, and Johnson 2020). For variation from unknown sources, Surrogate Variable Analysis (SVA) and Remove Unwanted Variation (RUV) are commonly used to generate surrogate variables, which help account for latent factors, reduce dependence, and stabilize error rate estimates in downstream differential expression analysis (Leek et al. 2012; Gagnon-Bartsch and Speed 2012).

Despite the success of ComBat, ComBat-seq, SVA, and RUV in their respective domains, the direct application of the aforementioned tools to DNA methylation data remains challenging. This is because DNA methylation data consist of methylation percentages, also known as β-values representing the proportion of methylated alleles at specific genomic loci. β-values are constrained within the range of 0 to 1, and the underlying distribution of β-values often deviates from a Gaussian distribution, exhibiting skewness and over-dispersion. To address this issue, β-values can be converted to M-values through a logit transformation prior to batch correction (Tian et al. 2017). Beyond these general strategies, several methods have been specifically developed for methylation data, including the two-stage RUVm, a variant of RUV, and BEclear (Maksimovic et al. 2015; Akulenko, Merl, and Helms 2016). However, the effectiveness of these approaches has yet to be systematically investigated.

Here we introduce ComBat-met, a method tailored specifically for adjusting batch effects in DNA methylation data. Building upon the principles of ComBat and ComBat-seq, our method employs a beta regression framework to account for the specific characteristics of β-values. ComBatmet estimates the parameters of a beta regression model, calculates a batch-free distribution, i.e., the expected distribution as if there were no batch effects in the data, based on the estimated model parameters, and adjusts the data by mapping the quantiles of the estimated distribution to its batch-free counterpart (Fig. 1). Besides the common cross-batch average, we also allow adjustment of β-values to a reference batch, where all batches are adjusted to the mean and precision of the reference (Zhang et al. 2018).

**Fig 1.**
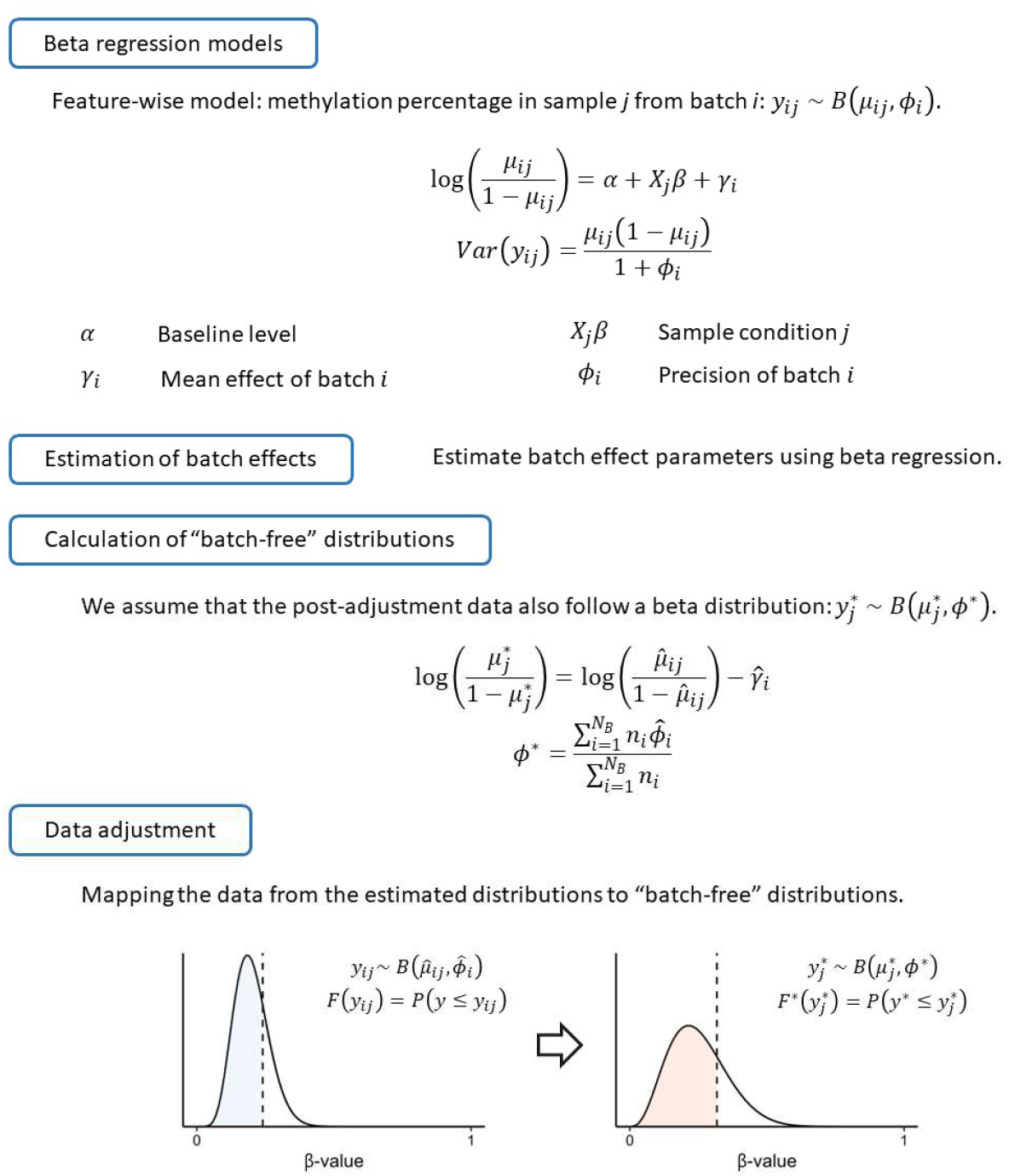
Diagram of the ComBat-met workflow. Model parameters are estimated using beta regression followed by the calculation of a batch-free distribution. A quantile-matching procedure is subsequently applied to map data from the estimated distribution to its batch-free counterpart.

In the rest of this paper, we describe the methodology of ComBat-met and demonstrate its performance through comprehensive benchmarking analyses and real-world applications. By utilizing simulated data, we demonstrated that ComBat-met followed by differential methylation analysis achieved superior statistical power compared to traditional approaches while correctly controlling the Type I error rate in nearly all cases. Furthermore, to illustrate its ability to recover biologically meaningful results and influence downstream analyses, we applied ComBatmet to batch-correct the methylation data in breast cancer patients pulled from The Cancer Genome Atlas (TCGA) (Cancer Genome Atlas 2012). Our results suggest that ComBat-met provides an effective solution for mitigating batch effects in methylation data, enabling more accurate and reliable downstream analyses.

## 2 Methods

### 2.1 Beta regression model

Let *y*_*ij*_ denote the β-value of a feature in sample *j* from batch *i*. A feature could be a site, a gene, or any molecular entity for which the methylation percentage could be defined. *y*_*ij*_ is assumed to follow a beta distribution, where *µ*_*ij*_ and *ϕ*_*i*_ denote the mean and precision of the distribution. The beta regression model is defined as:

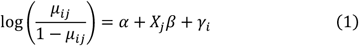

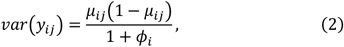

where *α, X*_*j*_, and *β* denote the common cross-batch average M-value (i.e., logit transformation of β-values), the covariate vector, and the corresponding regression coefficients, respectively. *γ*_*i*_ represents the batch-associated additive effect on the common cross-batch average M-value. To ensure identifiability, the sum of *γ*_*i*_ weighted by the sample size in each batch is constrained to zero (Johnson, Li, and Rabinovic 2007). Parameters are estimated via maximum likelihood estimation using the betareg() function in the betareg R package (Cribari-Neto and Zeileis 2010).

In the context of reference-based adjustment, *α* and *γ*_*i*_ in Eq. (1) represent the average M-value in the chosen reference batch and the additive difference between the reference batch and batch *i*, respectively. The rest of the notations remain unchanged.

### 2.2 Adjustment

To adjust the data, we take a quantile-matching approach similar to that described in ComBat-seq (Fig. 1) (Zhang, Parmigiani, and Johnson 2020). We assume that the adjusted data follow a beta distribution, i.e., 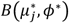. Once the regression model is fitted to the data, we calculate the parameters for the batch-free distributions, 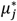 and *ϕ*^*^, as follows:

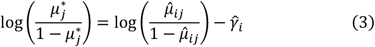

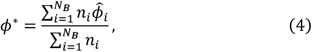

where *N*_B_ is the number of batches, *n*_*i*_ is the number of samples in batch *i*, and 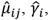, and 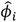 represent the maximum likelihood estimates of *µ*_*ij*_, *γ*_*i*_, and *ϕ*_*i*_ in Eq. (1) and Eq. (2), respectively. The adjusted data 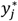 are then calculated by matching the quantile of the original data *y*_*ij*_ on the estimated distribution 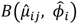 to its quantile counterpart on the batch-free distribution, i.e., 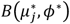. Specifically, each *y*_*ij*_ is mapped to its counterpart 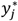 such that the cumulative distribution of 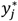, i.e., 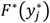 is closest to that of *y*_*ij*_, i.e., *F*(*y*_*ij*_) (Fig. 1). For reference-based adjustment, 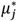 and *ϕ*^*^ are the maximum likelihood estimates of the mean and precision in the reference batch, respectively.

### 2.3 Simulation

The simulation was set up as follows. A total of 1000 features were generated with a balanced design involving two biological conditions and two batches across 20 samples (N = 20). Without loss of generality, we assumed that 100 out of 1000 features were truly differentially methylated, and the methylation percentage of these features was higher under condition 2 than under condition 1 by 10%. Data simulation was implemented using the dataSim() function in the methylKit R package, yielding 1000 × 20 matrices of coverages (i.e., total numbers of bases) and numbers of methylated bases (Akalin et al. 2012). β-values were calculated as the ratios of numbers of methylated bases to coverages for each feature.

All features were affected by batch. Both the mean and precision of batch effects were allowed to vary. The methylation percentage in one batch was assumed to be either higher or lower than that in the other batch by 0%, 2%, 5%, or 10%. The precision in one batch was set to a baseline value equal to 10, and the precision in the other batch was set to be 1-, 2-, 5-, or 10-fold of the baseline value. The remaining simulation parameters were resolved to their default values in dataSim() (Akalin et al. 2012).

The simulation was repeated 1000 times. The proportion of significant features among those that were and were not truly differentially methylated were counted towards true positive rates (TPRs) and false positive rates (FPRs), respectively. Median TPRs and FPRs over the 1000 repeated simulations were calculated under each parameter setting.

### 2.4 Other approaches

To validate ComBat-met, we compared its performance with several additional batch correction workflows: naïve ComBat, the “one-step” approach, M-value ComBat, SVA, RUVm, ComBat-biseq, and BEclear. Naïve Combat, in which beta-values are directly batch-corrected without first transforming them to M-values, is included in Section 3.1 solely to demonstrate the importance of using appropriate model assumptions.

The “one-step” approach, M-value ComBat, SVA, and RUVm all required that β-values be logit-transformed to M-values, which were implicitly assumed to be normally distributed, prior to batch correction. The “one-step” approach included the batch variable directly as a covariate in the differential linear model. M-value ComBat adjusted M-values for batch effects through ComBat (Johnson, Li, and Rabinovic 2007). SVA computed a surrogate variable to be included as a covariate in the downstream differential analysis without reference to the given batch information (Leek et al. 2012). Similarly, RUVm extended the RUV framework by leveraging control features to estimate and adjust for unwanted variation in methylation data (Maksimovic et al. 2015). BEclear, in contrast, applies latent factor models to identify batch-associated variations and replace them with values reconstructed from neighboring data entries (Akulenko, Merl, and Helms 2016).

To accommodate bisulfite sequencing data, we also developed ComBat-biseq, a method which retains the integer nature of count data. Unlike ComBat-met, ComBat-biseq initiates batch correction by fitting beta-binomial regression models to methylated cytosine counts. The remaining steps, however, are akin to ComBat-met (see Supplementary Fig. 1 for details). Beta-binomial regression, as well as the distribution and quantile functions of the beta-binomial distribution were implemented using the aod and updog R packages (Lesnoff and Lancelot 2012; Gerard et al. 2018). Differential methylation analysis of the adjusted bisulfite sequencing data was implemented using a likelihood ratio test (Park et al. 2014).

### 2.5 Application to real-world experimental data

Level 3 Illumina 450K methylation data (calculated β-values mapped to the genome) for tumor and adjacent normal tissues in breast cancer (BRCA) patients were downloaded from TCGA using the TCGAbiolinks R package (Colaprico et al. 2016; Mounir et al. 2019). To evaluate precision correction, we retained only batches containing at least two samples for batch adjustment. Furthermore, we focused exclusively on tumor samples in the luminal B subtype cohort for demonstration and visualization purposes. Our analysis included a total of 11 batches spanning 96 adjacent normal tissue samples and 19 batches spanning 135 tumor tissue samples. Similar to a previous study, tumor and adjacent normal tissue samples were analyzed separately to account for the larger inter-patient heterogeneity in tumor tissues (Supplementary Figs. 2 and 3) (Akulenko, Merl, and Helms 2016).

The probe-level β-value matrices were adjusted to the reference batch”A12R”, which was present in both tumor and normal samples, using the ComBat-met and M-value ComBat workflows. To corroborate our findings, batch correction was also performed at the gene level by aggregating probe-level data to the nearest genes, following a previous study (Robinson, McCarthy, and Smyth 2010; Hansen and Aryee 2012; Akulenko, Merl, and Helms 2016). Additional details on this process are provided in the Supplementary Methods.

To demonstrate the impact of batch correction on downstream machine learning, a neural network was trained to classify samples as normal or tumor based on methylation percentages. A minimalistic architecture was used, consisting of an input layer (3 randomly selected genes), two hidden layers (the first with 16 nodes and the second with 4 nodes), and an output layer. Gene-level data were used instead of probe-level data to avoid the high correlation between neighboring probes, which can lead to redundant features. Random feature selection ensured that the model’s performance was not biased by any gene set, and this process was repeated 50 times. The model was trained on data both before and after ComBat-met adjustment, and the performance was compared using accuracy metrics across 50 iterations.

## 3 Results

### 3.1 Using appropriate model assumptions for β-values

To demonstrate the importance of applying appropriate statistical models, we simulated an example of a β-value matrix with a balanced design, consisting of 20 samples (N = 20) pulled from two batches and two conditions (Fig. 2). While naïve ComBat, which applies ComBat directly to the β-value matrix, reduces cross-batch variations, doing so also causes adjusted β-values to exceed the maximum plausible limit of one in samples #12 and #13, posing challenges for data interpretation and downstream analysis (Fig. 2). This problem arises because ComBat operates based on a Gaussian model (Johnson, Li, and Rabinovic 2007). Similar to count data, β-values exhibit skewness and over-dispersion that cannot be captured by a Gaussian distribution.

**Fig 2.**
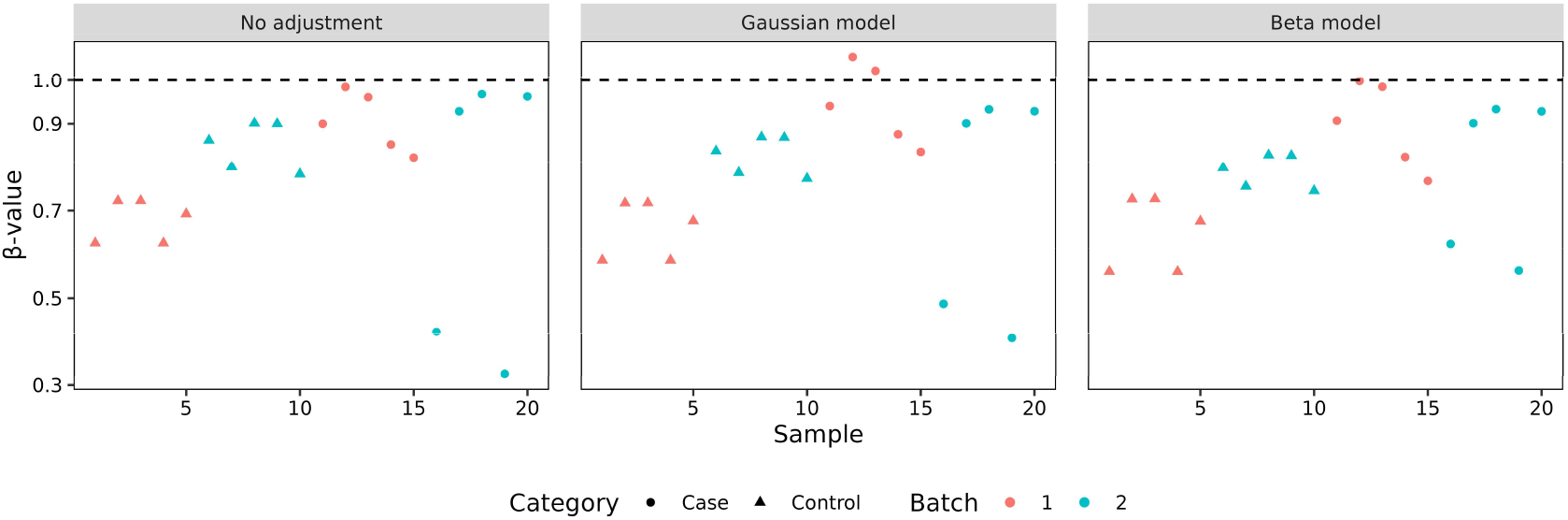
Example of problems caused by Gaussian model-based batch correction in DNA methylation data. A β-value matrix of 20 samples was simulated with a balanced design involving two batches and two categories. The left, middle, and right panels represent the unadjusted, Gaussian model-adjusted, and beta model-adjusted β-values across 20 samples, respectively. Batches are shown by color, and categories are shown by shape. Dashed lines represent the maximum plausible β-value equal to one.

Unlike ComBat, ComBat-met assumes that β-values follow a beta distribution, a statistical model much more suitable for the behavior of proportions and percentages. As such, it not only imposes a naturally confined range of (0, 1) on the adjusted β-values but also renders parameter estimation more robust to outliers, yielding the smallest cross-batch variations post-adjustment (Fig. 2).

### 3.2 ComBat-met demonstrates balanced control of power and false positive rates

We evaluated the FPR of ComBat-met using simulated DNA methylation data. In our simulations, the cross-batch differences in the mean and precision of the methylation percentage were allowed to vary. The resulting data were batch-corrected either using ComBat-met, M-value ComBat, ComBat-biseq, SVA, RUVm, or BEclear prior to differential methylation analysis, or by directly including the batch variable in the analysis (i.e., “one-step” approach). Details of each method are provided in Section 2.4.

The comparison of different batch correction methods is provided in Fig. 3. ComBat-met, M-value ComBat, and the “one-step” approach demonstrated close control of the FPR around 0.05 in most cases (Fig. 3). Although ComBat-met and M-value ComBat exceeded the threshold of 0.05 for small cross-batch differences in mean and precision (e.g., a 2% mean difference and a 2-fold precision difference) (Fig. 3), a slightly inflated FPR could generally be tolerated for discovery studies.

**Fig 3.**
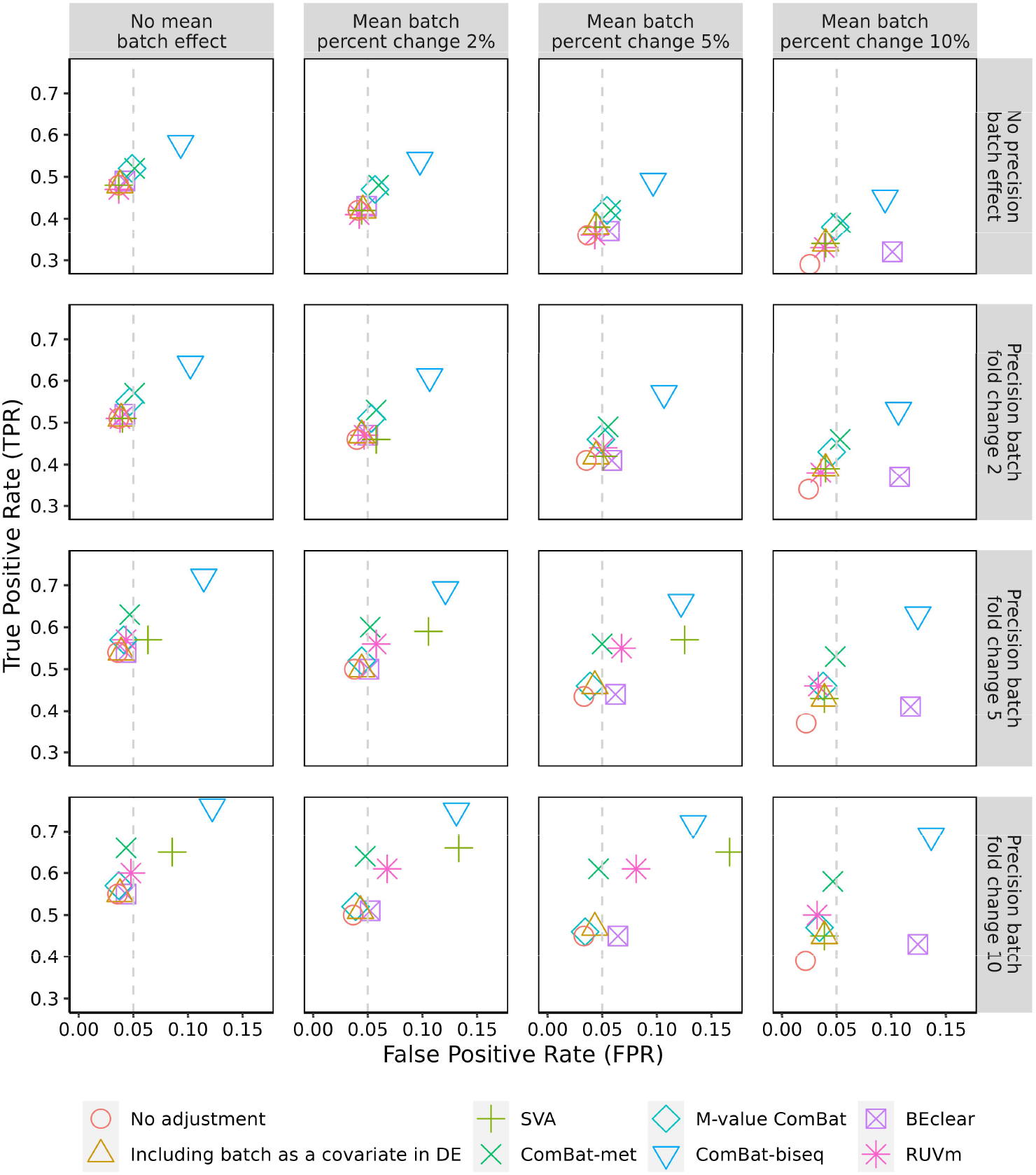
Median true positive rates and false positive rates calculated based on simulation data. The cross-batch mean difference in methylation percentage was set to 0%, 2%, 5%, or 10%. The precision of the batch effect was set to have a 1-, 2-, 5-, or 10-fold change. The simulation was repeated 1000 times. Methods are shown by color. Detailed information about each method is provided in Section 2.4.

In contrast to the aforementioned methods, ComBat-biseq consistently exhibited severe inflations of FPRs (Fig. 3). This behavior was most likely attributed to the small sample size, leading to breakdowns in the asymptotic argument of the likelihood ratio test downstream ComBat-biseq. Additional simulations supported this hypothesis, as the same workflow maintained much more precise control of the FPR for a larger sample size (N = 100) (Supplementary Fig. 4). BEclear showed markedly inflated FPRs for scenarios involving a 10% cross-batch mean difference, underscoring its limitations in handling large batch effects (Fig. 3). On the other hand, SVA and RUVm showed unstable performance across different parameter settings, with inflated FPRs in some cases and reduced TPRs in others (Fig. 3). This variability was expected, considering that SVA and RUVm, unlike ComBat-derived methods, are not optimized for scenarios where technical variations originate from identifiable sources.

The TPRs of these methods were evaluated based on the 10% features that were truly differentially methylated. All batch correction methods yielded higher TPRs than no adjustment, especially notable for a 10% cross-batch difference in the mean methylation percentage (Fig. 3).

Among different methods, ComBat-met and M-value ComBat achieved similar TPRs under circumstances with zero or a 2-fold cross-batch precision difference (Fig. 3). However, for scenarios involving a 5- or 10-fold precision change, ComBat-met demonstrated a markedly increased power to detect methylation changes compared to M-value ComBat, without compromising the FPRs (Fig. 3). This highlights the superior performance of ComBat-met, as large precision differences typically occur when combining data from heterogeneous batches, such as those from different profiling platforms or studies. While ComBat-biseq achieved even higher TPRs than ComBat-met, its limited ability to control FPRs significantly restricts its suitability for small sample sizes (Fig. 3).

### 3.3 ComBat-met improves batch adjustment and enhances machine learning performance

To demonstrate the ability of ComBat-met to recover biological insights, we applied our method to batch-correct the TCGA dataset described in Subsection 2.5. In adjacent normal tissues, samples from the same batches were clustered together in the unadjusted data (Supplementary Fig. 5). Both M-value ComBat and ComBat-met effectively mitigated batch effects, as evidenced by the absence of batch-related sample separation in the adjusted data (Supplementary Fig. 5). Analysis of explained variation further suggested that batch-related variance was markedly reduced in the data adjusted by ComBat-met compared to M-value ComBat (Fig. 4a). While the tumor samples were seemingly less batch-affected due to larger inter-patient heterogeneity, quantitative analysis also revealed the greatest reduction of batch-explained variance in the data adjusted by ComBatmet, similar to adjacent normal samples (Supplementary Fig. 6, Fig. 4a). Gene-level analysis post aggregation of β-values to the nearest genes revealed similar patterns, lending additional credibility to the efficacy of ComBat-met (Supplementary Figs. 7-9). Interestingly, applying ComBatmet with parameter shrinkage led to incomplete reduction of batch effects (Supplementary Fig. 10; see Supplementary Methods for details of parameter shrinkage).

**Fig 4.**
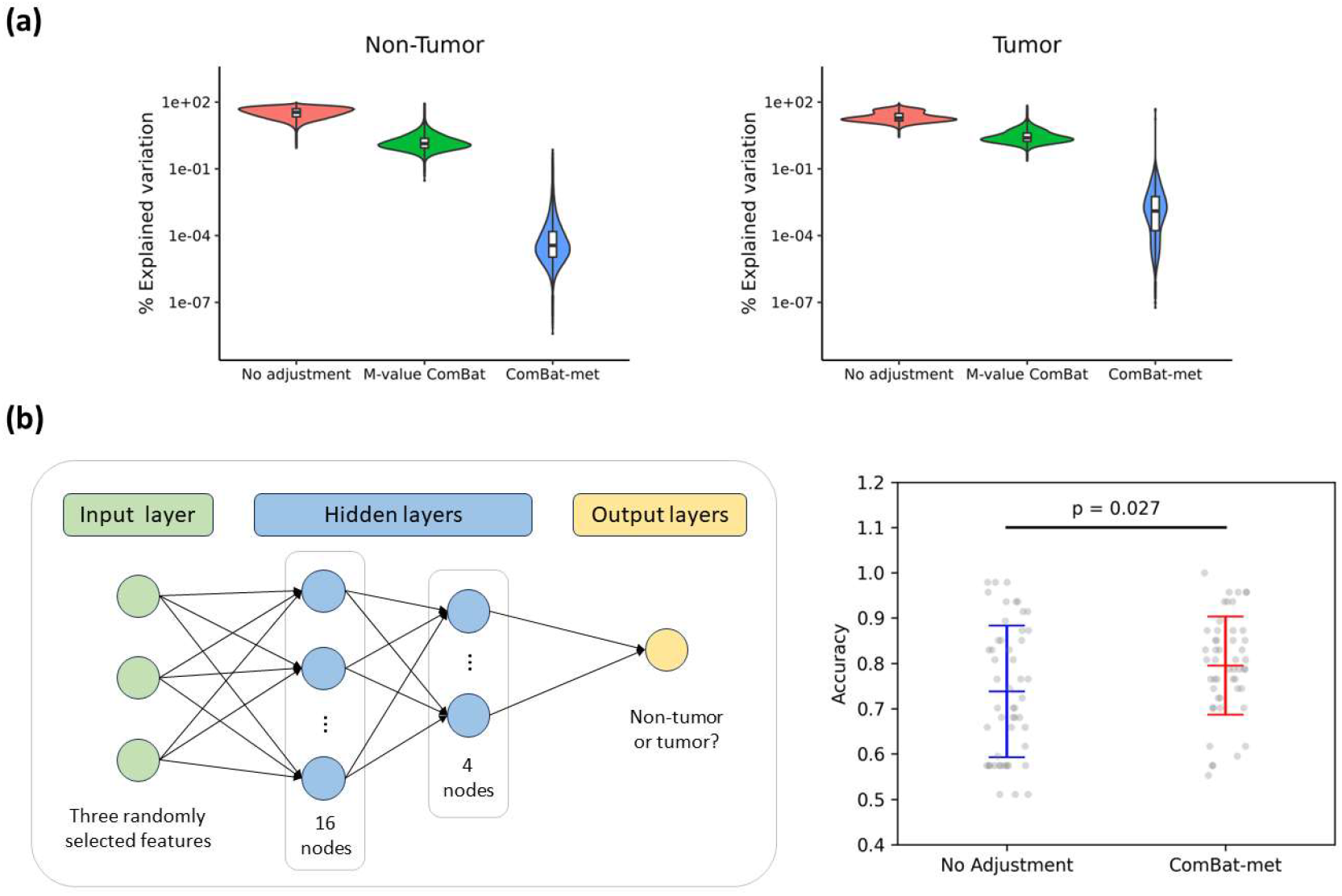
Application of ComBat-met to remove batch effects in the TCGA data. (a) Percent of variation explained by batch in the unadjusted probe-level data, probe-level data adjusted by M-value ComBat, and probe-level data adjusted by ComBat-met in tumor and adjacent normal samples. (b) Neural network architecture and a jitter plot showing improved classification accuracy with ComBat-met compared to no batch adjustment. Each gray dot represents the result from one of 50 iterations, with error bars indicating the mean +/-standard deviation (SD). The p-value was calculated using a two-sample t-test.

The impact of batch correction on downstream classification tasks was evaluated using a neural network trained to distinguish between normal and tumor samples based on methylation percentages. Models trained on batch-adjusted data outperformed those trained on uncorrected data, with a mean accuracy of 0.80 (SD: 0.11) for ComBat-met compared to 0.74 (SD: 0.14) for no adjustment (Fig. 4b). This improvement highlights the importance of effective batch correction in preserving biological signal while minimizing unwanted variation.

## 4 Discussion

In this work, we have developed ComBat-met, which adjusts batch effects in DNA methylation data by employing a beta regression framework. Our work not only underscores the importance of batch correction in methylation studies but also highlights the superior performance of ComBat-met through simulations and real-world applications. Compared to the widely adopted M-value ComBat approach, ComBat-met demonstrates more effective removal of batch effects and markedly improved sensitivity to detect methylation changes for scenarios involving large cross-batch precision differences, without sacrificing specificity.

ComBat-met is an extension of ComBat and ComBat-seq (Johnson, Li, and Rabinovic 2007; Zhang, Parmigiani, and Johnson 2020). Despite the differences in their probabilistic assumptions, all these methods adopt a regression framework to model the means and precision of batch effects followed by an adjustment step, which is in essence, a quantile-mapping procedure. Similar to ComBat, ComBat-met also provides users the option to adjust β-values to not only the common cross-batch average but also any existing batch as the reference (Zhang et al. 2018). In large consortium projects, datasets are typically generated sequentially (Subramanian et al. 2017; Wang and Novick 2023), so batch correction would always need to be reapplied in the event of incoming new datasets if the correction method depends on all datasets in hand. Our reference-based strategy allows new data to be adjusted without impacting the previous results, facilitating data integration across sequential batches.

While ComBat-met brings major improvement to batch correction for methylation studies, we acknowledge several limitations of our approach. First, ComBat-met runs more slowly than M-value ComBat. Unlike linear regression, beta regression lacks a closed-form solution and therefore requires iterative methods for maximum likelihood estimation. Parallel computing may be employed to expedite the regression step in ComBatmet. Second, ComBat-met is designed specifically to address batch effects stemming from identifiable sources. In the presence of unknown sources of variation, SVA or RUV should be applied instead (Leek et al. 2012; Gagnon-Bartsch and Speed 2012).

Additionally, ComBat employs an empirical Bayes framework to shrink batch effect parameters (Johnson, Li, and Rabinovic 2007). This approach borrows information across features, making the adjustment robust for data with small sample sizes or outliers. Inspired by the non-parametric empirical Bayes method in ComBat (Johnson, Li, and Rabinovic 2007), we provide a similar option in ComBat-met, enabling users the freedom to shrink parameters at will. However, in the context of beta regression, parameter shrinkage has been shown to cause under-correction of batch effects (Supplementary Fig. 10). Our observations are consistent with ComBat-seq, which also under-corrects when coupled to parameter shrinkage. Similar to the negative binomial distribution, beta distributions can accommodate a wide range of skewness, conferring more robustness to outliers and making parameter shrinkage unnecessary (Zhang, Parmigiani, and Johnson 2020). Furthermore, it is noteworthy that nonparametric empirical Bayes, which requires extensive Monte-Carlo sampling, becomes extremely time-consuming for large DNA methylomes even if only a subset of features are sampled. While we discourage the use of parameter shrinkage in ComBat-met, it will be interesting to further explore the impact of outlying β-values and refine our method to better accommodate outliers in methylation data.

## 5 Conclusions

In conclusion, ComBat-met emerges as a powerful batch correction method for capturing the unique characteristics of DNA methylation data. Our work demonstrates the great potential of this approach in facilitating data integration and accelerating biological discoveries.

## Code availability

ComBat-met, M-value ComBat, and ComBat-biseq, along with the scripts used to generate the simulated data and to analyse the simulated data and the data from TCGA, are available as an R package at: https://github.com/JmWangBio/ComBatMet.

