## supplement for "ComBat-met: Adjusting Batch Effects in DNA Methylation Data"

Junmin Wang[1,*]

[1] *Data Sciences and Quantitative Biology, Discovery Sciences, Biopharmaceuticals R&D, AstraZeneca, Waltham, Massachusetts*

### Supplementary Information

- Supplementary Methods
- Supplementary Figures
- Supplementary References

**Supplementary Methods**

*Aggregation of β-values*

Aggregation of β-values was conducted following the workflow described in a previous study (Akulenko, Merl, and Helms 2016). First, we removed all sites/probes with missing β-values, as principal component analysis requires complete datasets. Next, the remaining probes were mapped to genes and filtered based on their distance to the respective transcription start sites (TSS) (Robinson, McCarthy, and Smyth 2010; Hansen and Aryee 2012). Depending on the strand direction, only probes within 2000 base pairs up- or downstream the respective TSS were retained for further analysis. Subsequently, the β-values of the probes were averaged by the nearest gene, resulting in a total of 4895 and 4639 gene-β-value pairs in normal and tumor tissues, respectively. The resulting β-value matrices were then separately adjusted to the reference batch “A12R”, which was present in both tumor and normal samples, using the ComBat-met and M-value ComBat workflows.

*Parameter Estimation with Shrinkage*

Inspired by ComBat and ComBat-seq, we provide a similar non-parametric empirical Bayes method to shrink parameters in ComBat-met (Johnson, Li, and Rabinovic 2007; Zhang, Parmigiani, and Johnson 2020). Let $y_{sij}$ denote the β-value of feature $s$ in sample $j$ from batch $i$. $y_{sij}$ is assumed to follow a beta distribution, where $\mu_{sij}$ and $\phi_{si}$ denote the mean and precision of the distribution. The beta regression model is defined as:

$$\log\left(\frac{\mu_{sij}}{1-\mu_{sij}}\right) = \alpha_s + X_j\beta_s + \gamma_{si}$$

$$var(y_{sij}) = \frac{\mu_{sij}(1-\mu_{sij})}{1+\phi_{si}},$$

where $\alpha_s$, $\beta_s$, $\gamma_{si}$, and $\phi_{si}$ are defined the same as in the main text for each feature $s$. In the empirical Bayes framework, the estimated values of $\gamma_{si}$ and $\ln[\phi_{si}]$ are adjusted to the means of the posterior distributions. Specifically, the empirical Bayes estimates of $\gamma_{si}$ and $\ln[\phi_{si}]$ are calculated as the weighted average across features:

$$\hat{\gamma}_{si}^{*} = \frac{\sum_{k\neq s}^{K} \omega_{ki}\hat{\gamma}_{ki}}{\sum_{k\neq s}^{K} \omega_{ki}}$$

$$\ln[\hat{\phi}_{si}^{*}] = \frac{\sum_{k\neq s}^{K} \omega_{ki}\ln[\hat{\phi}_{ki}]}{\sum_{k\neq s}^{K} \omega_{ki}},$$

where $\hat{\gamma}_{ki}$ and $\ln[\hat{\phi}_{ki}]$ denote the maximum likelihood estimates of $\gamma_{ki}$ and $\ln[\phi_{ki}]$, respectively. $K$ is the total number of features, or a subset of features selected to approximate the posterior distribution. The weights $\omega_{ki}$ are defined as the beta likelihood functions:

$$\omega_{ki} = \prod_{j=1}^{n_i} d(y_{sij}|\hat{\gamma}_{ki}, \hat{\phi}_{ki}),$$

where $d$ denotes the density function of the beta distribution, and $n_i$ is the number of samples in batch $i$. Parameters for batch-free distributions, $\mu_{sj}^{*}$ and $\phi_s^{*}$, are calculated as:

$$\log\left(\frac{\mu_{sj}^{*}}{1-\mu_{sj}^{*}}\right) = \log\left(\frac{\hat{\mu}_{sij}}{1-\hat{\mu}_{sij}}\right) - \hat{\gamma}_{si}^{*}$$

$$\phi_s^{*} = \frac{\sum_{i=1}^{N_B} n_i\hat{\phi}_{si}^{*}}{\sum_{i=1}^{N_B} n_i}.$$

### Supplementary Figures

**Beta-binomial regression models**

Feature-wise model: methylated count in sample *j* from batch *i*: $y_{ij} \sim BetaBin(T_j, \mu_{ij}, \phi_i)$.

$$\log\left(\frac{\mu_{ij}}{1-\mu_{ij}}\right) = \alpha + X_j\beta + \gamma_i$$

$$Var(y_{ij}) = T_j\mu_{ij}(1-\mu_{ij})\left[1 + \frac{T_j - 1}{1+\phi_i}\right]$$

| | | | |
|---|---|---|---|
| $\alpha$ | Baseline level | $X_j\beta$ | Sample condition *j* |
| $\gamma_i$ | Mean effect of batch $i$ | $\phi_i$ | Precision of batch $i$ |
| $T_j$ | Total count in sample *j* | | |

**Estimation of batch effects**

Estimate batch effect parameters using beta-binomial regression.

**Calculation of "batch-free" distributions**

We assume that the post-adjustment data also follow a beta-binomial distribution: $y_j^* \sim BetaBin(T_j, \mu_j^*, \phi^*)$.

$$\log\left(\frac{\mu_j^*}{1-\mu_j^*}\right) = \log\left(\frac{\hat{\mu}_{ij}}{1-\hat{\mu}_{ij}}\right) - \hat{\gamma}_i$$

$$\phi^* = \frac{\sum_{i=1}^{N_B} n_i\hat{\phi}_i}{\sum_{i=1}^{N_B} n_i}$$

**Data adjustment**

Mapping the data from the estimated distributions to "batch-free" distributions.

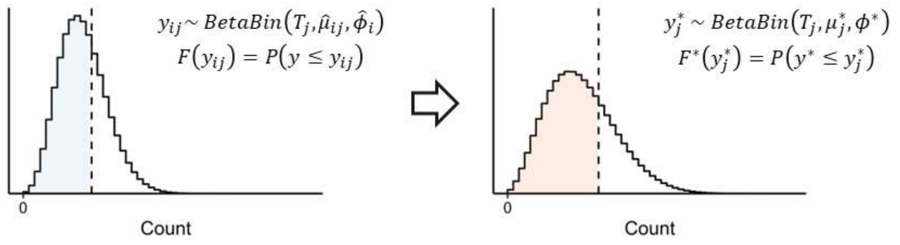


**Supplementary Fig. 1. Diagram of the ComBat-biseq workflow.**

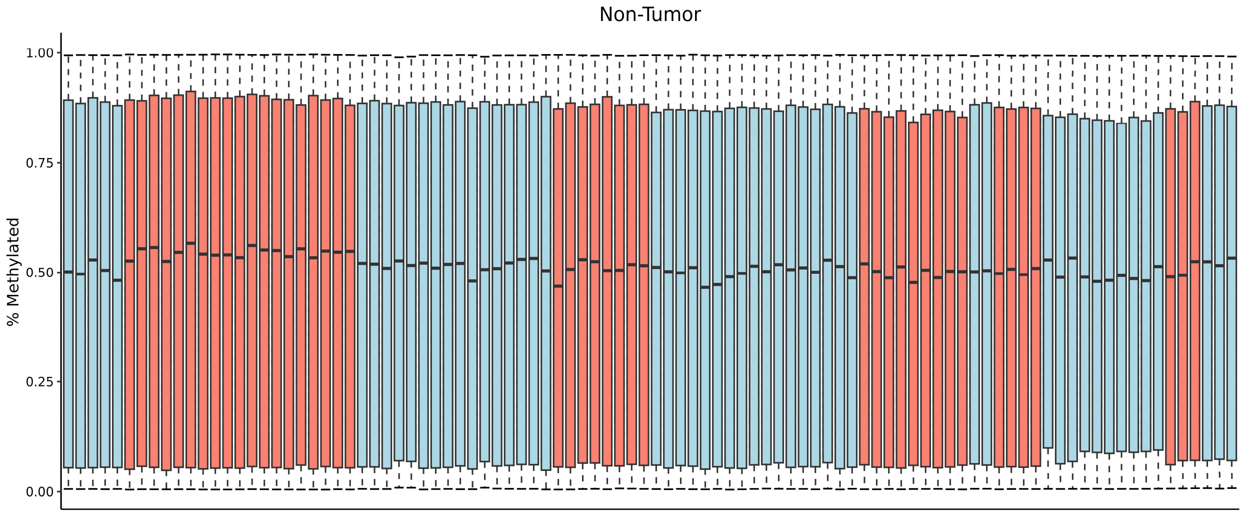


**Supplementary Fig. 2. Box plot illustrating the distribution of β-values per sample in the adjacent normal tissues of breast cancer patients.** Batches are shown by color.

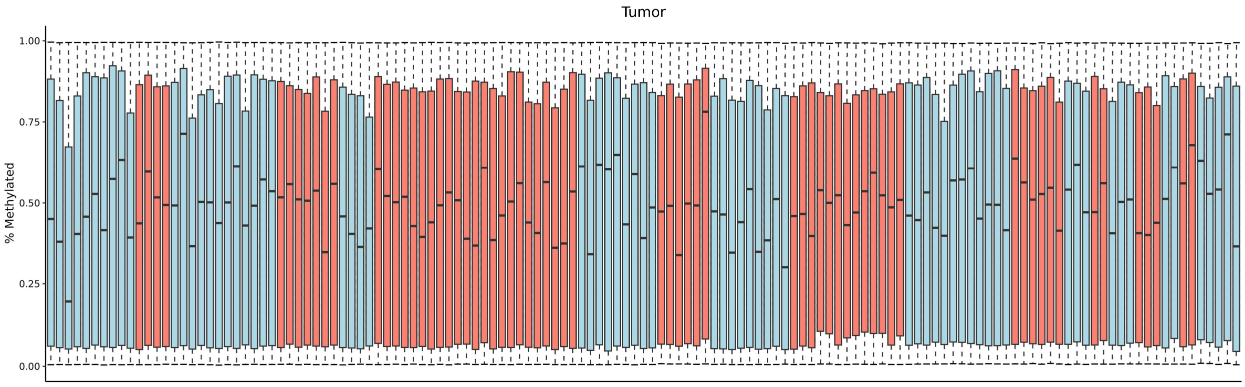


**Supplementary Fig. 3. Box plot illustrating the distribution of β-values per sample in the tumor tissues of breast cancer patients.** Batches are shown by color.

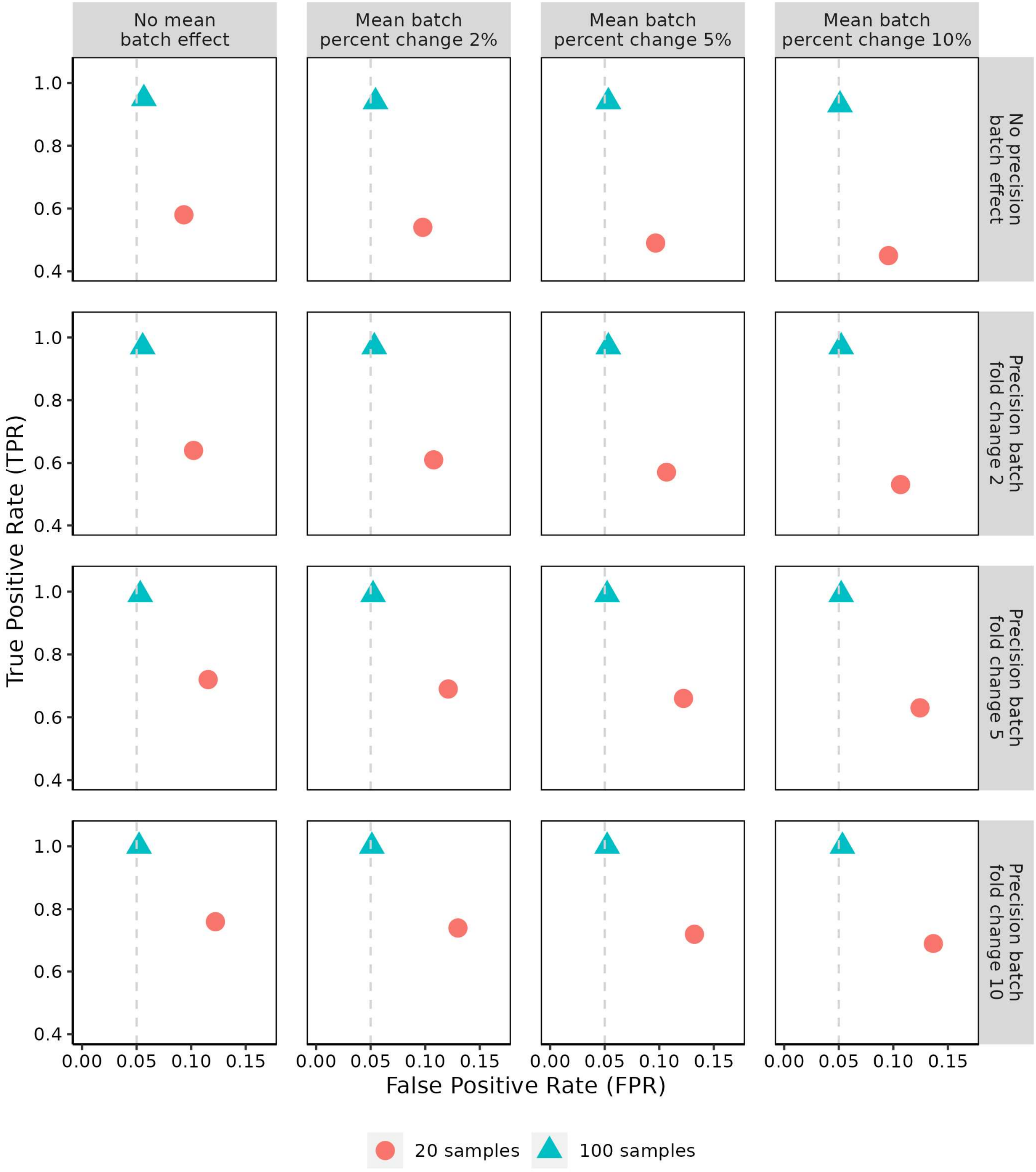


**Supplementary Fig. 4. Median true positive rates and false positive rates of ComBat-biseq based on simulation data.** Either 20 or 100 samples were simulated. The cross-batch mean difference in methylation percentage was set to 0%, 2%, 5%, or 10%. The precision of the batch effect was set to have a 1-, 2-, 5-, or 10-fold change. The simulation was repeated 1000 times.

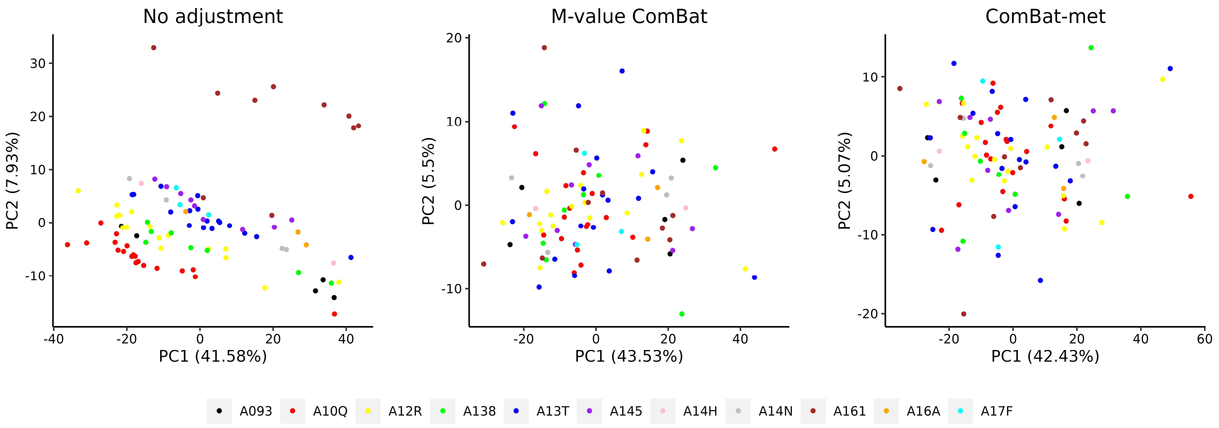


**Supplementary Fig. 5. PCA plots illustrating the separation of adjacent normal tissue samples in the unadjusted probe-level data, probe-level data adjusted by M-value ComBat, and probe-level data adjusted by ComBat-met.** Batches are shown by color.

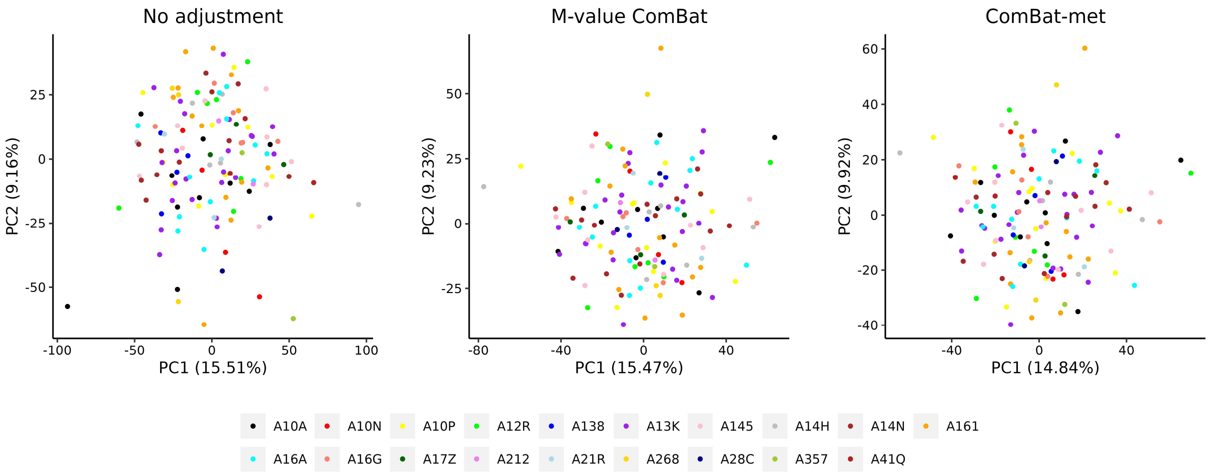


**Supplementary Fig. 6. PCA plots illustrating the separation of tumor tissue samples in the unadjusted probe-level data, probe-level data adjusted by M-value ComBat, and probe-level data adjusted by ComBat-met.** Batches are shown by color.

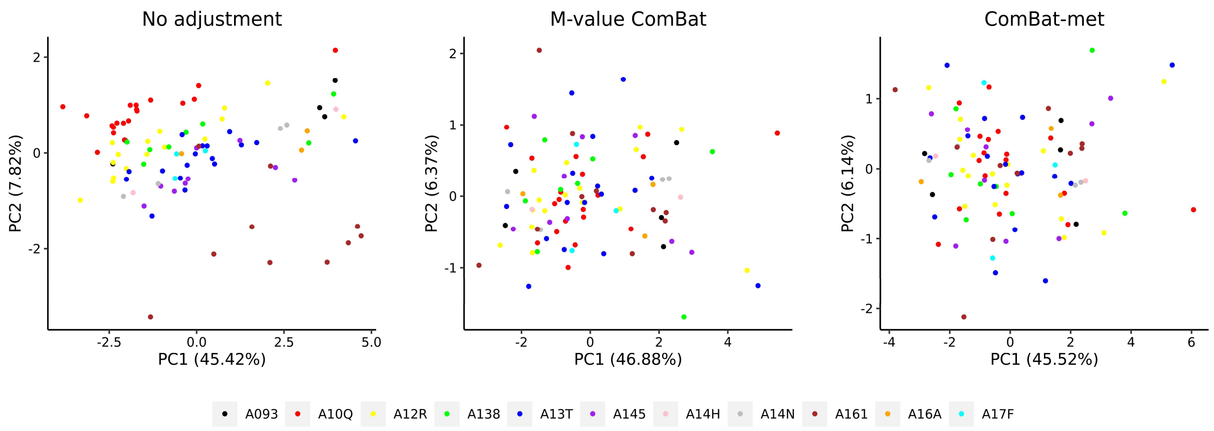


**Supplementary Fig. 7. PCA plots illustrating the separation of adjacent normal tissue samples in the unadjusted gene-level data, gene-level data adjusted by M-value ComBat, and gene-level data adjusted by ComBat-met.** Batches are shown by color.

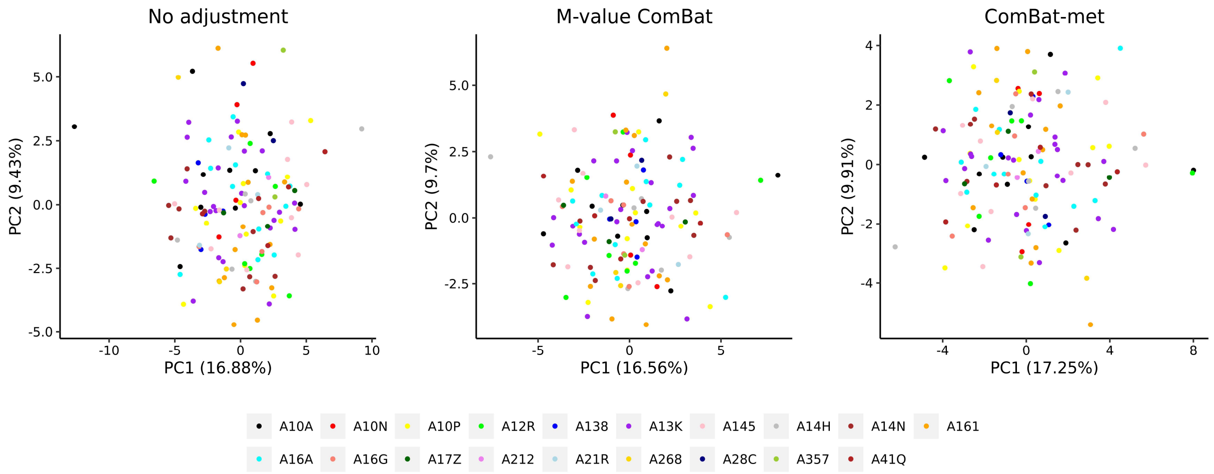


**Supplementary Fig. 8. PCA plots illustrating the separation of tumor tissue samples in the unadjusted gene-level data, gene-level data adjusted by M-value ComBat, and gene-level data adjusted by ComBat-met.** Batches are shown by color.

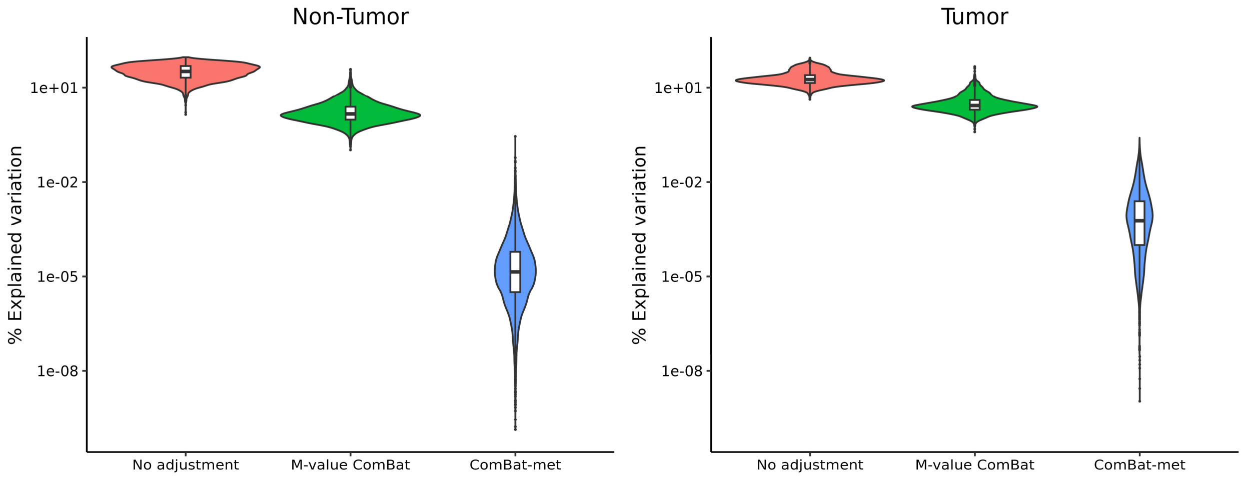


**Supplementary Fig. 9. Percent of variation explained by batch in the unadjusted gene-level data, gene-level data adjusted by M-value ComBat, and gene-level data adjusted by ComBat-met in tumor and adjacent normal samples.**

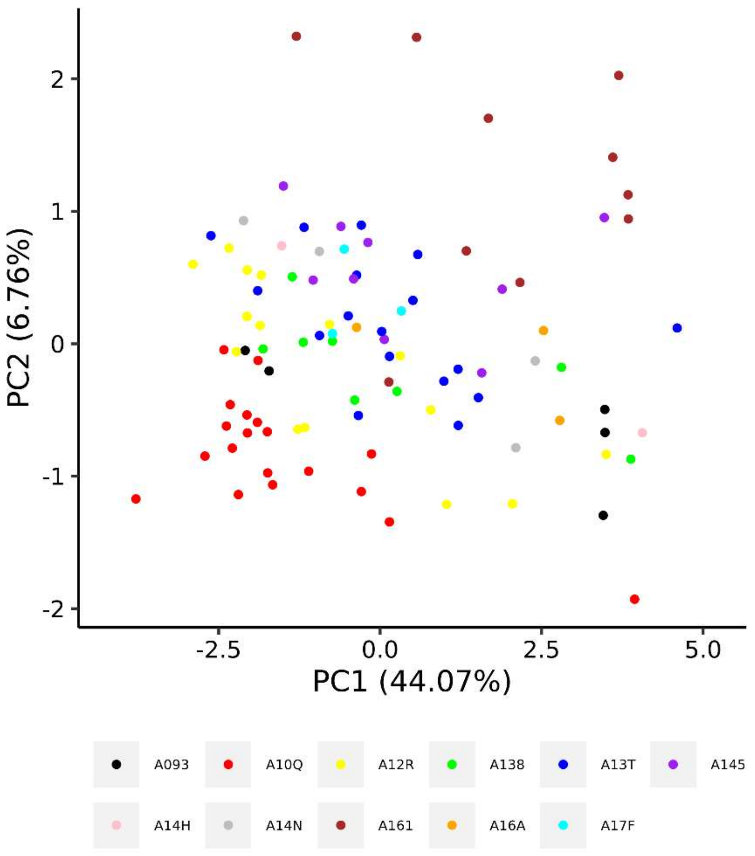


**Supplementary Fig. 10. PCA plot illustrating the separation of adjacent normal tissue samples in the data adjusted by ComBat-met followed by parameter shrinkage.** Batches are shown by color.
